# Erratic Maternal Care Induces Avoidant-Like Attachment Deficits in a Mouse Model of Early Life Adversity

**DOI:** 10.1101/2025.06.13.659607

**Authors:** Zoë A. MacDowell Kaswan, Christian Bowers, Ivan Teplyakov, Jose Munoz-Martin, Sahabuddin Ahmed, Lily Kaffman, Lauryn Giuliano, Marcelo O. Dietrich, Arie Kaffman

## Abstract

Attachment theory offers an important clinical framework for understanding and treating negative effects of early life adversity. Attachment styles emerge during critical periods of development in response to caregivers’ ability to consistently meet their offspring’s needs. Attachment styles are classified as secure or insecure (anxious, avoidant, or disorganized), with rates of insecure attachment rising in high-risk populations and correlating with a plethora of negative health outcomes throughout life. Despite its importance, little is known about the neural basis of attachment. Work in rats has demonstrated that limited bedding and nesting (LB) impairs maternal care and produces abnormal maternal attachment linked to increased pup corticosterone. However, the effects of LB on attachment-like behavior have not been investigated in mice where additional genetic and molecular tools are available. Furthermore, no group has utilized home-cage monitoring to link abnormal maternal care with deficits in attachment-like behavior. Using home-cage monitoring, we confirmed a robust increase in maternal fragmentation among LB dams. Abnormal maternal care was correlated with elevated corticosterone levels on post-natal day seven (P7) and a stunted growth trajectory that persisted later in life. LB did not alter maternal buffering at P8 or maternal preference at P18, indicating that certain attachment-like behaviors remain unaffected despite exposure to high levels of erratic maternal care. However, LB pups vocalized less in response to maternal separation at P8, did not readily approach their dam at P13, and exhibited higher anxiety-like behavior at P18, suggesting that LB induces avoidant-like attachment deficits in mice.

**Significance Statement:** The impoverished conditions of limited bedding and nesting (LB) cause erratic maternal care and elevated corticosterone levels in rat and mouse pups. The increase in corticosterone levels causes attachment-like deficits in rat pups; however, it remains unclear whether similar deficits are observed in mice, where additional genomic and molecular tools are available. Using continuous home-cage monitoring, we confirmed a substantial increase in erratic maternal care and elevated corticosterone levels in 7-day-old mouse pups. LB mouse pups exhibited attachment-like deficits in some, but not all, tests, underscoring the robustness of this evolutionarily conserved bond. Despite some similarities, the attachment abnormalities observed in mice differed from previous reports in rats, paving the way for in-depth mechanistic studies in mice.

## Introduction

According to attachment theory, reciprocal interactions between a caregiver and their offspring during a critical developmental period, establish a strong emotional bond that is essential for survival and normal development (Bowlby, 1978; Izaki et al., 2024). A typical attachment sequence begins with distress vocalizations from the offspring in response to homeostatic perturbations, leading to proximity-seeking behaviors that bring the offspring and the caregiver closer together to alleviate distress. The repeated successful completion of this sequence creates a secure base from which the offspring can explore their environment (Hornor, 2019; Sullivan and Opendak, 2021; Izaki et al., 2024)

Using the Strange Situation paradigm as a standardized procedure for observing infant responses to separation and reunification from the caregiver, Ainsworth and colleagues identified four attachment styles: secure, anxious, avoidant, and disorganized, with anxious, avoidant, and disorganized styles sometimes grouped together as insecure attachment (Hornor, 2019; Gregory et al., 2020). Secure attachment occurs when caregivers consistently meet the child’s physical and emotional needs, which is manifested by the caregiver’s capacity to effectively console and reduce distress upon reunification. Anxiously attached children exhibit excessive distress when the caregiver leaves and are difficult to console upon reunion. In contrast, avoidantly attached children display minimal distress when the caregiver departs and may avoid or even express anger towards the caregiver during reunion. Children with disorganized attachment demonstrate a combination of insecure and avoidant behaviors (Hornor, 2019; Gregory et al., 2020). Rates of disorganized attachment can be as high as 35% among children who have experienced severe neglect or maltreatment (Wright and Edginton, 2016) and are correlated with significant psychiatric and somatization disorders later in life (Hornor, 2019; Gregory et al., 2020).

Attachment-based randomized controlled trials have been shown to increase rates of secure attachment and to improve psychiatric comorbidities in high-risk populations (Wright and Edginton, 2016; Diamond et al., 2019; Gregory et al., 2020; Hepworth et al., 2020). Additionally, these trials have been effective in improving medical comorbidities (Hornor, 2019; Gregory et al., 2020), including reducing inflammation (Ross et al., 2021; Londono Tobon et al., 2023), and decreasing stunting (Noviana et al., 2023), further demonstrating the clinical utility of this approach.

Despite the significance of attachment formation during development, our understanding of its underlying biology remains limited. This includes the impact of attachment on brain circuits that regulate mood and cognition, as well as more general processes such as immune function and growth (Izaki et al., 2024). The evolutionarily conserved nature of attachment suggests that rodent models can offer valuable insights into these questions (Sullivan and Opendak, 2021). Studies in rats have shown that erratic maternal care induced by limited bedding and nesting material (LB) leads to elevated pup corticosterone levels that are directly responsible for abnormal amygdala activation and deficits in proximity-seeking behavior (Raineki et al., 2019). However, to the best of our knowledge, the effects of LB on maternal attachment have not yet been investigated in mice, where additional genetic tools could be utilized to study the neurobiology of attachment behavior.

The goals of the present study were to establish a reproducible method for measuring maternal behavior and to develop a longitudinal series of tests for assessing maternal attachment in mouse pups using the LB paradigm. We found that maternal movement and fragmentation during the first week of life were highly quantifiable and reproducible using a home-cage monitoring system and present data suggesting that LB pups display avoidant-like attachment deficits. Together, this study provides a foundation to further investigate the mechanisms by which erratic maternal care disrupts attachment behavior in the mouse.

## Methods

### Animals

C57BL/6J (Jackson Laboratories stock #000664) mice were housed in standard Plexiglas cages and kept on a standard 12:12 h light-dark cycle (lights on at 07.00 AM) with constant temperature 20 ± 1 °C and humidity 43% ± 2, with food and water provided ad libitum. All studies were approved by the Institutional Animal Care and Use Committee (IACUC) at Yale University and were conducted in accordance with the recommendations of the NIH Guide for the Care and Use of Laboratory Animals.

### Limited Bedding (LB)

C57BL/6J mice were mated using a 3:1 female to male ratio in standard mouse Plexiglas cages layered with 500 cc of corncob bedding and a nestlet. Visibly pregnant females were placed individually in Noldus PhenoTyper cages for mice (Noldus Technology) approximately 2-4 days before birth and given 500 cc softcob bedding (Cat# 7087C, Inotiv Teklad), a nestlet, and 2-3 chow pellets on the floor of the cage. The PhenoTypers are a home cage monitoring system comprised of a 30 x 30 cm cage with side-mounted food and water, and a lid with built-in infrared lights and an infrared camera. The PhenoTypers are directly connected to Noldus EthoVision XT 17 (Noldus Technology) software for automated continuous mouse tracking. The nest region of each arena was assigned in EthoVision and beginning at birth, designated as postnatal day (P0), the dams’ total movement, time in the nest, and frequency of entrances/exists to the nest were recorded continuously until P2 (pre-assignment period, black rectangle in Fig 1A). At P2, litters were culled to 5-8 pups per litter and randomly assigned to either control (CTL) or LB conditions. CTL litters received 500 cc of softcob bedding, a nestlet, and 15 cc soiled bedding from the old cage. LB litters received 125 cc softcob bedding, no nestlet, and 15 cc soiled bedding from the old cage. Dam behavior was recorded using EthoVision until P7 (post-assignment period, red rectangle in Fig 1A). At P7, litters were transferred to standard mouse Plexiglas cages with CTL litters receiving 500 cc of corncob bedding and a nestlet, and LB litters receiving 125 cc corncob bedding and no nestlet as previously described (Ahmed et al., 2024; Islam et al., 2024). Bedding was changed at P14 and P21. Litters weaned at P26 and housed with 2-5 same-sex littermates in standard Plexiglass mouse cages containing 500 cc of corncob bedding, a nestlet, and 2-3 pellets of chow on the floor of the cage.

**Figure 1.**
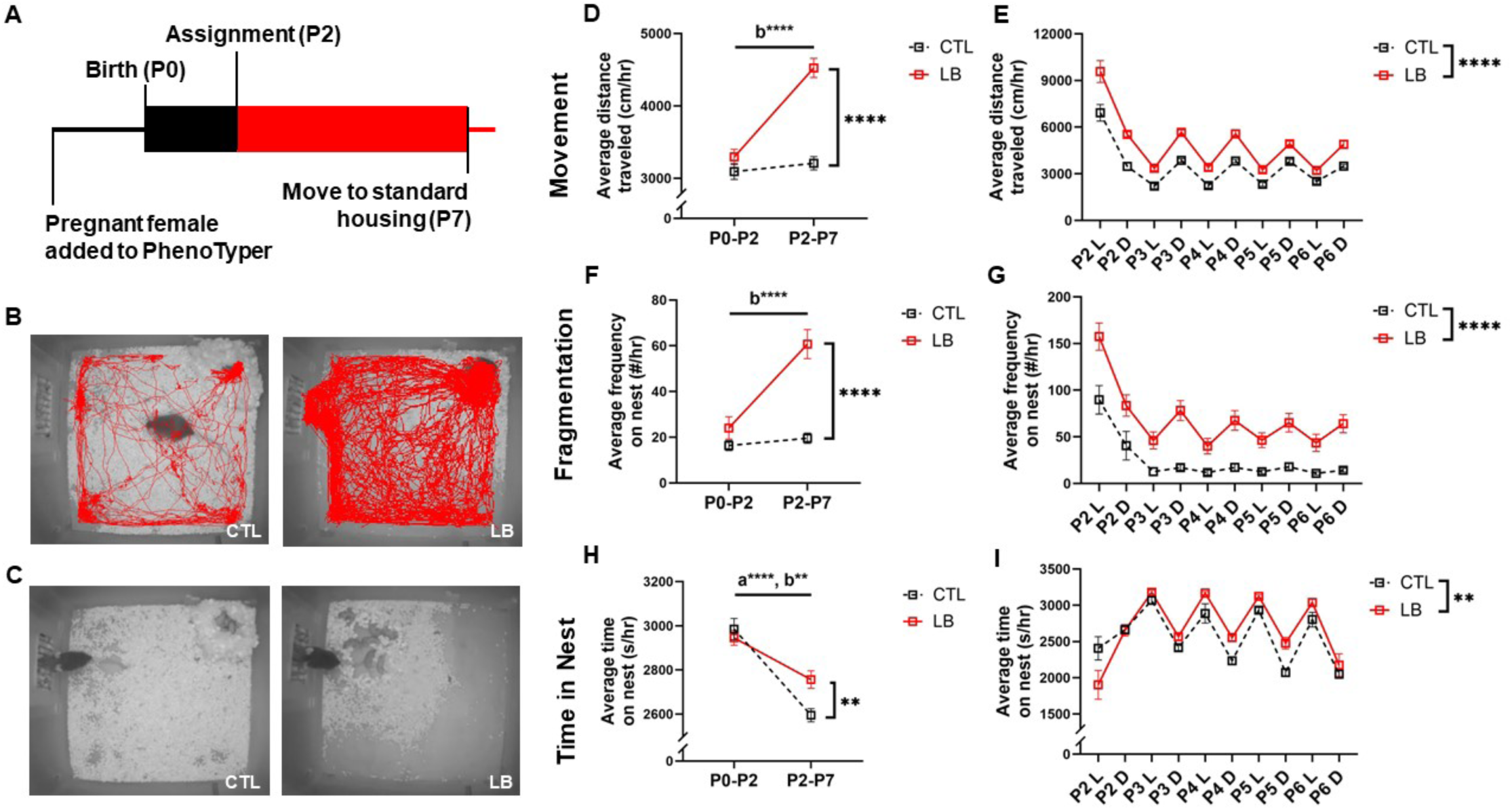
LB Increases Distance Traveled, Maternal Fragmentation, and Time in Nest. **(A)** PhenoTyper timeline. Black box indicates pre-assignment recording time, red box indicates post-assignment recording time. **(B)** Representative tracing of the dam’s movement from the 2nd to 3rd hour of the dark phase of P3. **(C)** Representative images of dams and their litters in the PhenoTypers during the dark phase of P3. **(D)** Average hourly distance traveled by the dam pre-(P0-P2) and post-assignment (P2-P7). Rearing x assignment Interaction: F(1, 18) = 35.42, P < 0.0001. Post-hoc analysis: CTL vs LB (P0-P2): P = 0.14; CTL vs LB (P2-P7): P < 0.0001; CTL (P0-P2) vs (P2-P7): P = 0.39; LB (P0-P2 vs P2-P7): P < 0.0001. **(E)** Average hourly distance traveled by the dam during the light (L) and dark (D) phases after condition assignment. Day: F(1.617, 124.9) = 125.7, P < 0.0001; Rearing: F(1, 78) = 88.59, P < 0.0001; Light-Dark cycle: F(1, 78) = 14.98, P = 0.0002; Day x Light-Dark interaction: F(4, 309) = 126.1, P < 0.0001; Rearing x Light-Dark: non-significant (NS); Day x Light-Dark x Rearing interaction: NS. Post-hoc analysis: P2 vs. all other days: P < 0.0001. **(F)** Average hourly maternal fragmentation pre-(P0-P2) and post-assignment (P2-P7). Rearing x Assignment Interaction: F(1, 18) = 26.72, P < 0.0001. Post-hoc analysis: CTL vs LB (P0-P2): P = 0.16; CTL vs LB (P2-P7): P < 0.0001; CTL (P0-P2) vs (P2-P7): P = 0.49; LB (P0-P2 vs P2-P7): P < 0.0001. **(G)** Average hourly maternal fragmentation during the light (L) and dark (D) phases after condition assignment. Day: F 1.727, 133.4) = 44.55, P < 0.0001; Rearing: F (1, 78) = 69.06, P < 0.0001; Light-Dark cycle: NS; Day x Rearing interaction: NS; Rearing x Light-Dark: F(4, 309) = 19.45, P < 0.0001; Day x Light-Dark x Rearing interaction: NS. Post-hoc analysis: P2 vs. all other days: P < 0.0001. **(H)** Average hourly time on nest pre-(P0-2) and post-assignment (P2-7). Rearing assignment interaction: F(1, 38) = 6.84, P=0.013. Post-hoc analysis: CTL vs LB (P0-P2): P = 0.48; CTL vs LB (P2-P7): P = 0.0040; CTL (P0-P2) vs (P2-P7): P < 0.0001; LB (P0-P2 vs P2-P7): P = 0.0015. **(I)** Average hourly time on nest during the light (L) and dark (D) phases after condition assignment. Day: F(2.596, 199.9) = 14.78, P < 0.0001; Rearing: F(1,78) = 8.08, P = 0.0057, Light-Dark cycle: F(1, 78) = 102.6, P < 0.0001, Day x Rearing: F(4, 308) = 7.59, P < 0.0001; Day x Light-Dark: F(4, 308)=43.42, P<0.0001; Rearing x Light-Dark: NS; Day x Light-Dark x Rearing interaction: NS. N= 19-21 per rearing condition, 2 x 2 rmANOVA for D, F, H with a and b representing effects of pre-and post-assignments for CTL and LB, respectively. Three-way rmANOVA for E, G, and I. NS-non-significant, P > 0.05.

### Quantification of Maternal behavior

Separate detection settings were required for light and dark cycles. For cohort 1 (used for longitudinal behavior), videos of dam behavior were analyzed after recording. An observer watched EthoVision’s automated tracking and, when necessary, adjusted the detection settings for accuracy and moved the assigned nest zone if the dam moved her nest. For cohorts 2 and 3 (used for corticosterone and maternal affiliation, respectively), automated mouse tracking was performed in real time and detection settings were automatically switched between the room’s light and dark cycles. After data collection, videos were checked to ensure that the dam kept her nest in the pre-assigned area. If the dam moved her nest, the recording time was noted, assigned nest zone was moved in EthoVision, and the data were re-analyzed. Dams’ mean movement, cumulative times in Arena and nest zones, and frequency in the nest zone were automatically calculated by EthoVision in one-hour time bins. If the last time bin was incomplete, it was discarded. In some instances, EthoVision failed to detect the dam when she was immobile on the nest for long periods of time. Therefore, any time that the dam was not detected in the Arena, she was assumed to be on the nest.

### Blood Collection and Corticosterone ELISA (P7)

Blood was collected at 14:00-15:00 of P7 pups using rapid decapitation. To minimize stress, blood was collected in a separate room. Blood was first collected from the dam, followed by pups, which were individually transferred in a cup containing home-cage bedding. After allowing the blood to clot for 30 minutes at room temperature, samples were centrifuged for 5 minutes at 2400xg at room temperature. Serum was collected and stored at -80 °C until use. At least one male and one female pup per litter, of approximately average weight for that litter, for a total of 7-8 pups per sex per rearing condition were selected for corticosterone analysis. Serum corticosterone levels were measured in duplicates using ELISA kit (Cat. # K014-H1, Arbor Assays, Ann Arbor, MI), according to the manufacturer’s instructions, using the 100 μl format.

### Buffering of Ultrasonic Vocalizations (P8)

The extent to which a female’s presence buffered pups’ ultrasonic vocalizations was tested at P8 (i.e., one day after litters were transferred to standard Plexiglas cages, Fig 4A). Testing was done between 13:00-17:00 under standard room illumination (600 lux). During the isolation phase, pups were placed individually in an empty soundproofed standard Plexiglass cage and ultrasonic vocalizations (USVs) were recorded for 5min using UltraSoundGate condenser microphone CM 16 connected via an UltraSoundGate 416 USGH audio device (Avisoft Bioacoustics, Berlin, Germany). After 5 min of isolation, an anesthetized unrelated adult female (“aunt”), fed the same diet as the dam, was placed in physical contact with the pup for 5 minutes and USVs were recorded for an additional 5 min (“With Aunt” phase, Fig. 4B). Aunts were anesthetized using 10/0.1 mg/kg ketamine/xylazine mixture dissolved in water. At the completion of the recordings, pups were weighed and returned to the home and isolation chambers cleaned with 70% ethanol in between pups. Numbers of USVs were quantified using VocalMat software (Fonseca et al., 2021) and “buffering” was calculated using the following formula: % Buffering = (# USVs with aunt - # USVs in isolation) / # USVs in isolation * 100.

### Maternal Affiliation (P13)

At P13, pups were tested for proximity-seeking behavior to their anesthetized dam, termed “maternal affiliation.” Testing was done between 13:00-18:00 under standard room illumination (600 lux). Dams were anesthetized with urethane (1.5 mg/kg) and placed in a standardized Plexiglass mouse cage layered with 100 cc clean corncob bedding mixed with 30 cc of dirty home-cage bedding. Pups were then individually placed at one end of the arena and allowed five minutes to freely explore the cage. Pup behavior was recorded using an overhead HD Pro Webcam C920 (Logitech) and videos were analyzed using EthoVision XT 17 (Noldus Technology) to determine the total time spent at the dam’s ventral or dorsal side (summed to calculate total interaction time) latency to approach, and mean distance between pup and dam (calculated as mean distance between pup and closer of either the dorsal or ventral zone). For binary approach versus no approach analysis, a pup was considered to have approached if it entered either the dorsal or ventral zones.

### Open Field and Maternal Preference Tests (P18)

P18 pups were tested in the open field and maternal preference tests between 13:00-17:00 under standard room illumination (approx. 600 lux). In the first phase, the pup was given 5 minutes to explore the empty arena (20 x 42 cm). This served both as an open field test and habituation to the arena. The pup was then weighed, individually marked, and returned to the home cage. Immediately following this habituation, the dam and a novel object were placed under wire pencil cups at opposite ends of the arena. The pup was placed in the center of the arena and given 5 minutes to freely investigate (Fig 6B). Location of the dam and novel object were switched between litters to reduce bias. Both phases of the test were recorded using an overhead HD Pro Webcam C920 (Logitech) and analyzed using EthoVision XT 17 (Noldus Technology). For the open field test, total movement and percent time in center (a central region 1/2 the width and length of the total arena) were calculated. For the maternal preference test, total movement and the amount of time the pup spent in an ∼2 cm radius (“zone”) around either the pencil cup containing the dam or novel object was quantified. Preference index was calculated as: (time spent in dam’s zone) *100 / (time spent in dam’s zone + time spent in novel object’s zone). Thus, preference index > 50% indicates preference for the dam.

### Social Interaction (P33-34)

The social interaction test was performed in adolescent mice ages P33-34. Testing was performed between 13:00-17:00 under standard room illumination (600 lux). Experimental animals were allowed 10 minutes to explore the empty arena (20 x 42 cm). After 10 min of habituation, an age- and sex-matched stranger mouse was added to the arena (Fig 7B) and social exploration was recorded for 10 min using an overhead HD Pro Webcam C920 (Logitech). Latency to first interaction initiated by the experimental mouse and total time that the experimental mouse interacted with the stranger were quantified by an observed blind to rearing condition or sex.

### Experimental Design and Statistical Analysis

Data were graphed and statistical analyses were performed using GraphPad Prism 10 (GraphPad Software, La Jolla, California, USA). Maternal behavior was analyzed using a 2 x 2 repeated measures (rm) ANOVA, with rearing as a between-subjects variable and assignment as a within-subjects variable (Fig 1 D, F, H). Significant rearing by assignment interactions were followed by Fisher’s least significant difference (LSD) post-hoc analyses. A three-way rmANOVA was employed to assess the effects of rearing (between-subjects variable), day (within-subjects variable), and light-dark phase (within-subjects variable) on maternal behavior, with significant interactions followed by Tukey’s-HSD post-hoc analyses (Fig 1 E, G, I). Three-way rmANOVAs were also utilized to determine the effects of rearing, sex, and test phase (Fig 4C) or side preference (Fig 5G). Two-way ANOVAs were conducted to examine the effects of rearing and sex on weight at different ages (Fig 2A), corticosterone levels (Fig. 3B), maternal buffering (Fig 4D), maternal affiliation (Fig 5 D-F), exploratory behavior in the open field (Fig 6 C-D), maternal preference index (Fig 6 E-F), and social interaction (Fig 7 B-C). Pearson correlations were calculated to characterize the relationship between the dams’ behavior and the following variables: body weight (Fig 2 B-F), corticosterone levels at P8, maternal affiliation at P13, time spent in the center of the open field, and maternal preference at P18. Chi-square analysis was used to assess the effect of rearing on the approach vs. no approach analysis in the maternal affiliation test (Fig 5C).

**Figure 2.**
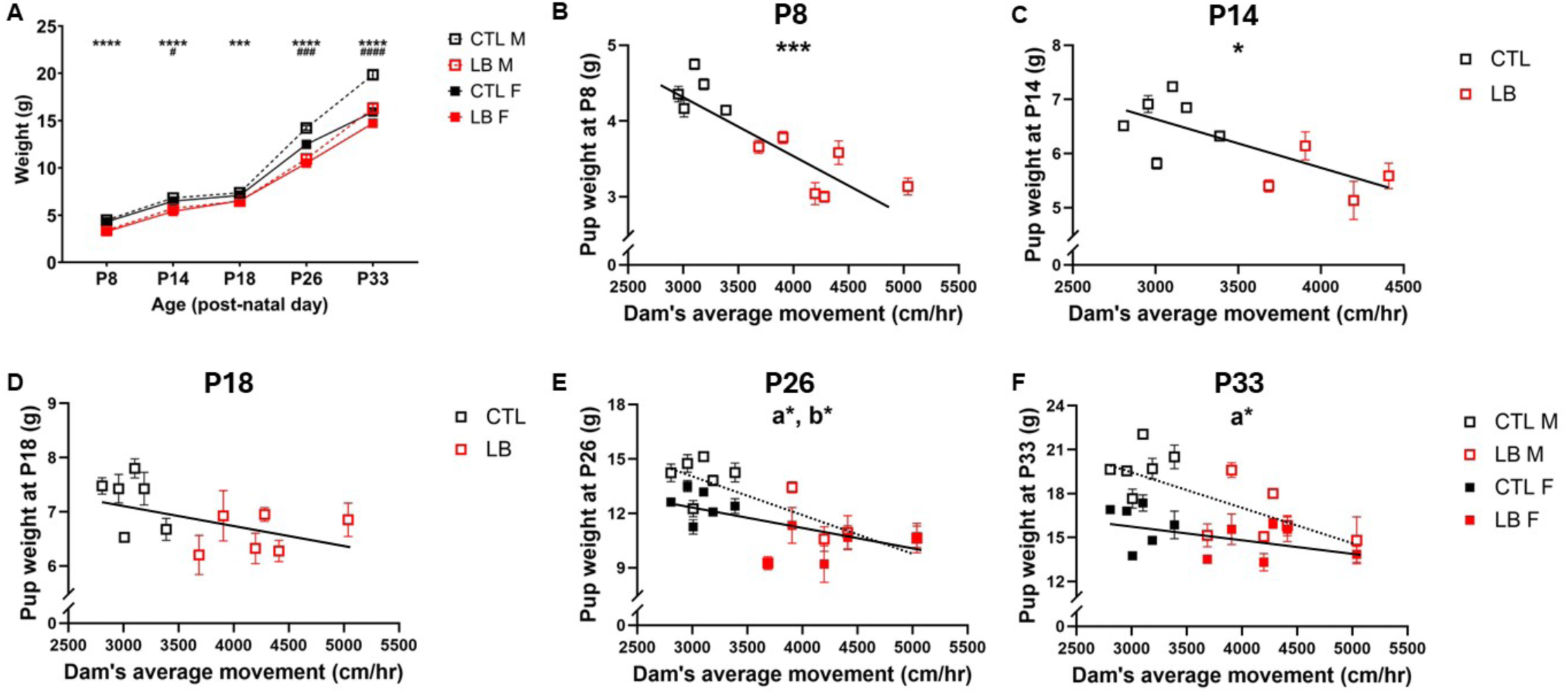
Dam’s Locomotor Activity During the First Week Negatively Correlates with Pup Bodyweight Throughout Development. **(A)** Effects of rearing and sex on pup weights at P8, P14, P18, P26, and P33; * indicates effect of rearing, ^#^ indicates effect of sex. P8: Rearing F (1, 72) = 153.4, P < 0.0001; Sex F (1, 72) = 3.68, P = 0.059; Interaction: NS. P14: Rearing F (1, 66) = 53.01, P < 0.0001; Sex F (1, 66) = 4.60, P = 0.036; Interaction: NS. P18: Rearing F (1, 44) = 14.75, P = 0.0004; Sex: NS; Interaction: NS. P26: Rearing F (1, 74) = 75.98, P < 0.0001; Sex F (1, 74) = 13.13, P = 0.0005; Interaction F (1, 74) = 4.514, P = 0.037. P33: Rearing F (1, 44) = 25.89, P < 0.0001; Sex F (1, 44) = 35.16, P < 0.0001; Interaction F (1, 44) = 6.42, P = 0.015. Correlation between pup body weight at P8 and dams’ average movement P2-7 at different ages. (**B**) P8: R^2^ = 0.76, P = 0.0005. **(C)** P14: R^2^ = 0.52, P = 0.019. **(D)** P18: R^2^ = 0.27, P = 0.08. **(E)** P26: Males: R^2^ = 0.49, P = 0.016, Females: R^2^ = 0.42, P = 0.030. **(F)** P33: Males: R^2^ = 0.50, P = 0.01, Females: R^2^ = 0.20, P = 0.14. Two-way ANOVA for each time points for A, with N = 4-6 litters, and N = 9-23 pups per rearing condition per time point. Pearson correlation for B-F for CTL (black) and LB (red) litters. Male and female pups were analyzed separately in E-F (male: open symbols, dotted line, labeled lower case ‘a’; female: closed symbols, solid line, labeled lower case ‘b’). NS-non-significant, P > 0.05.

**Figure 3:**
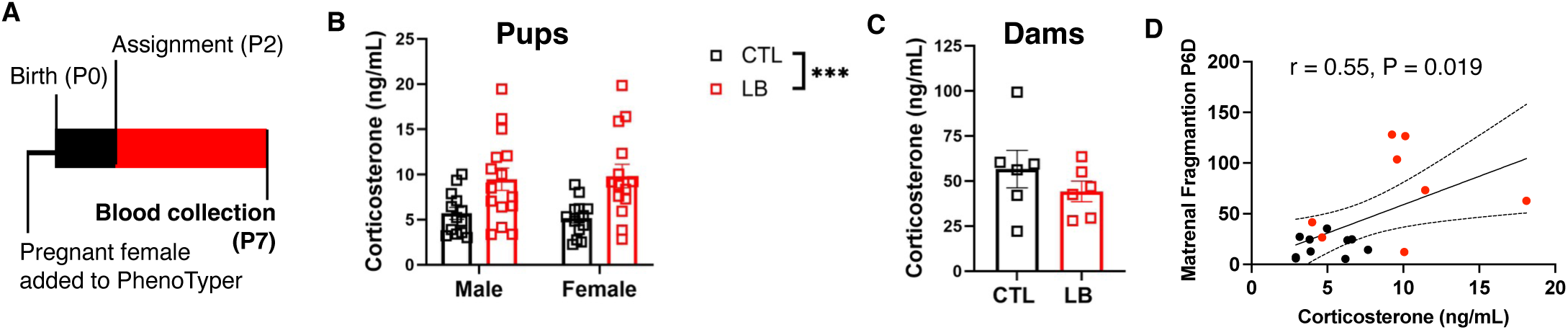
LB Increases Corticosterone Levels in P7 Pups. **(A)** Experimental timeline. **(B)** LB pups have higher, but more variable, corticosterone levels than CTL pups at P7; N=13-15 pups per sex per rearing condition from a total of 9 litters per condition. Two-way ANOVA, rearing: F(1,52) = 17.61, P = 0.0001; sex: NS; interaction: NS. **(C)** No difference between dams’ corticosterone levels at P7. Unpaired t-test, t(10) = 1.044, P = 0.32; N = 6 dams per condition. (**D**) Maternal fragmentation during the dark phase of P6 (prior to corticosterone assessment) correlated with the average corticosterone levels per liter obtained at P7, R^2^ = 0.30, P = 0.019, N = 2-4 pups per litter with nine litters per condition. NS-non-significant, P > 0.05.

**Figure 4:**
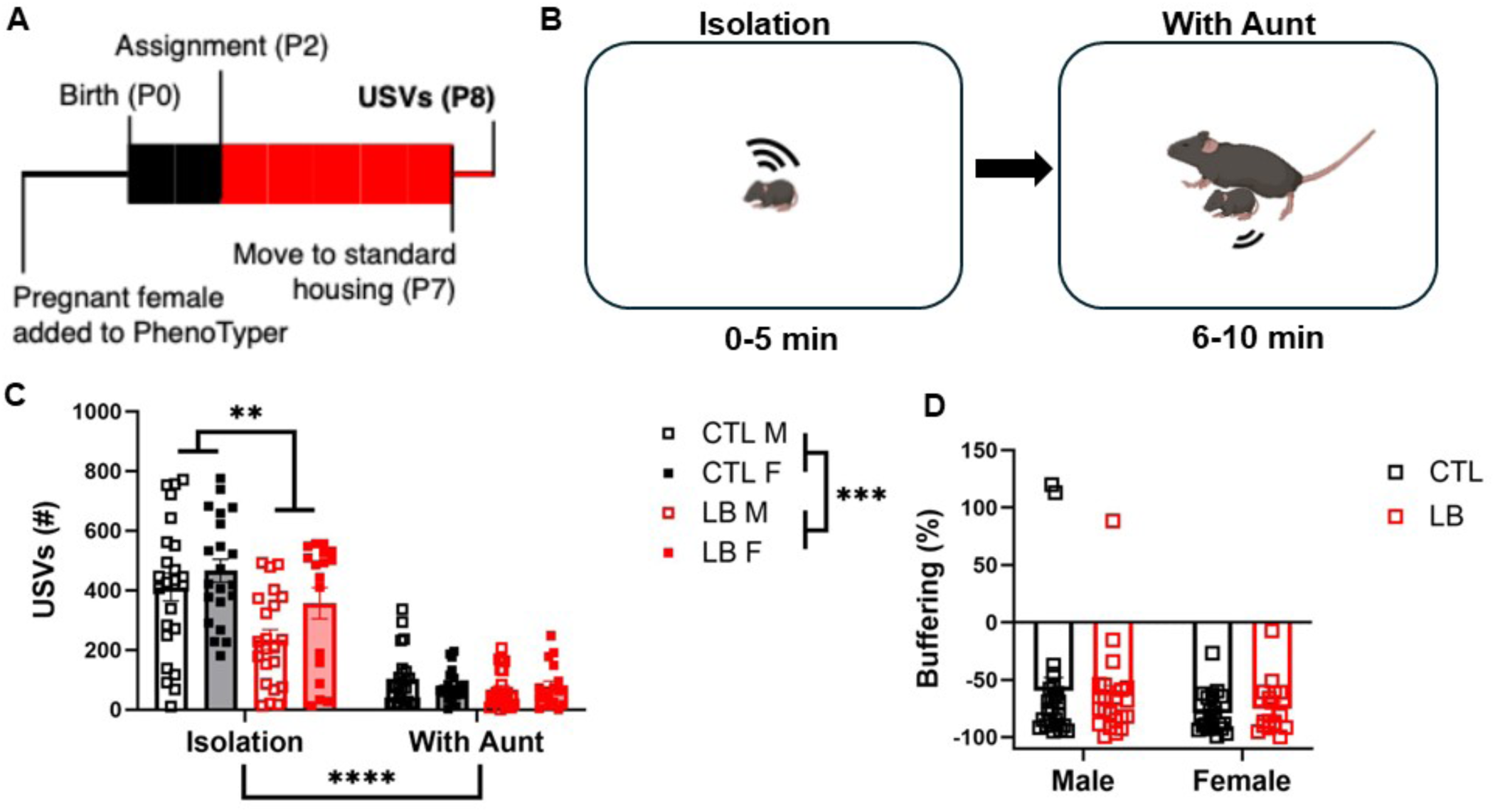
LB Pups Have Reduced Vocalization During Isolation but Do Not Differ in Buffering at P8. **(A)** Timeline. **(B)** Schematics of the test. **(C)** Total number of USVs during the two test phases. rmANOVA of isolation phase only, rearing: F(1, 72) = 8.276, P = 0.0053; sex: F(1, 72) = 3.68, P = 0.059; interaction: NS. rmANOVA of reunion phase only: rearing, sex and interaction are all NS **(D)** Percent change in vocalizations between phases (buffering). 2 x 2 ANOVA, rearing: NS; Sex: F(1, 80) = 3.34, P = 0.071; Interaction: NS. N=5-6 litters and N=17-24 pups per group. NS-non-significant, P > 0.05.

**Figure 5:**
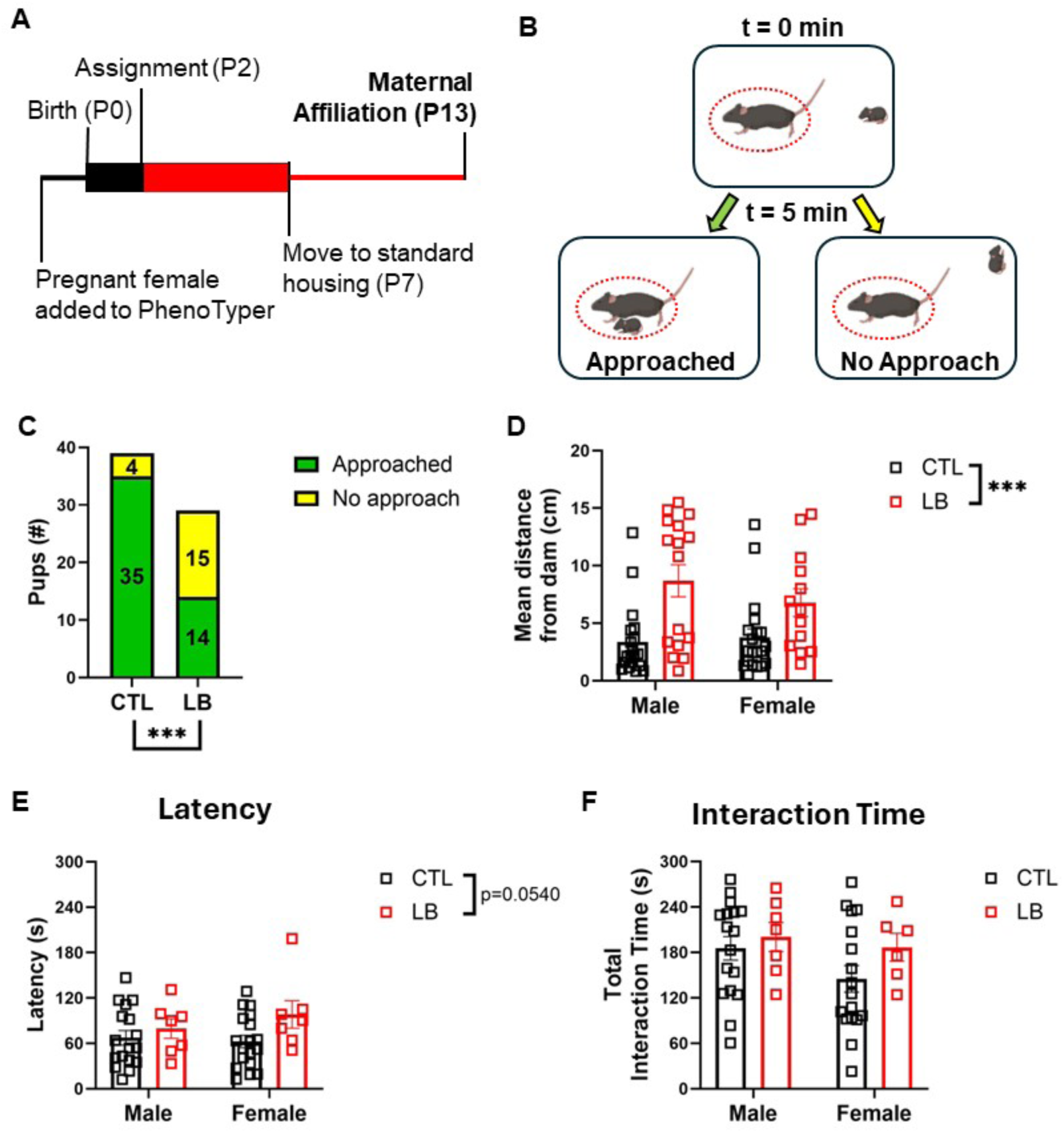
LB Impairs Approach Behavior in P13 Pups. **(A)** Timeline. **(B)** Experimental setup. Dotted red line indicates interaction zone. **(C)** 35 out of 39 (90%) of CTL pups approached the dam vs. 14 out of 29 (48%) of LB pups: X^2^ (1, 14.2) = 3.77, P = 0.0002. **(D**) Average distance from the dam. Rearing: F (1, 64) = 17.11, P = 0.001; Sex: NS; Interaction: NS. **(E)** Latency to approach the dam among pups that did so. Rearing: F(1, 45) = 3.92, P=0.054; Sex: NS; Interaction: NS. **(F) T**otal interaction time among pups that approached the dam. Rearing, sex and interaction were all NS. 2-way ANOVA in D, E, and F. N = 13-20 per sex and rearing condition (total of 39 CTL and 29 LB pups), from N = 6 CTL and 5 LB litters. NS-non-significant, P > 0.05.

**Figure 6:**
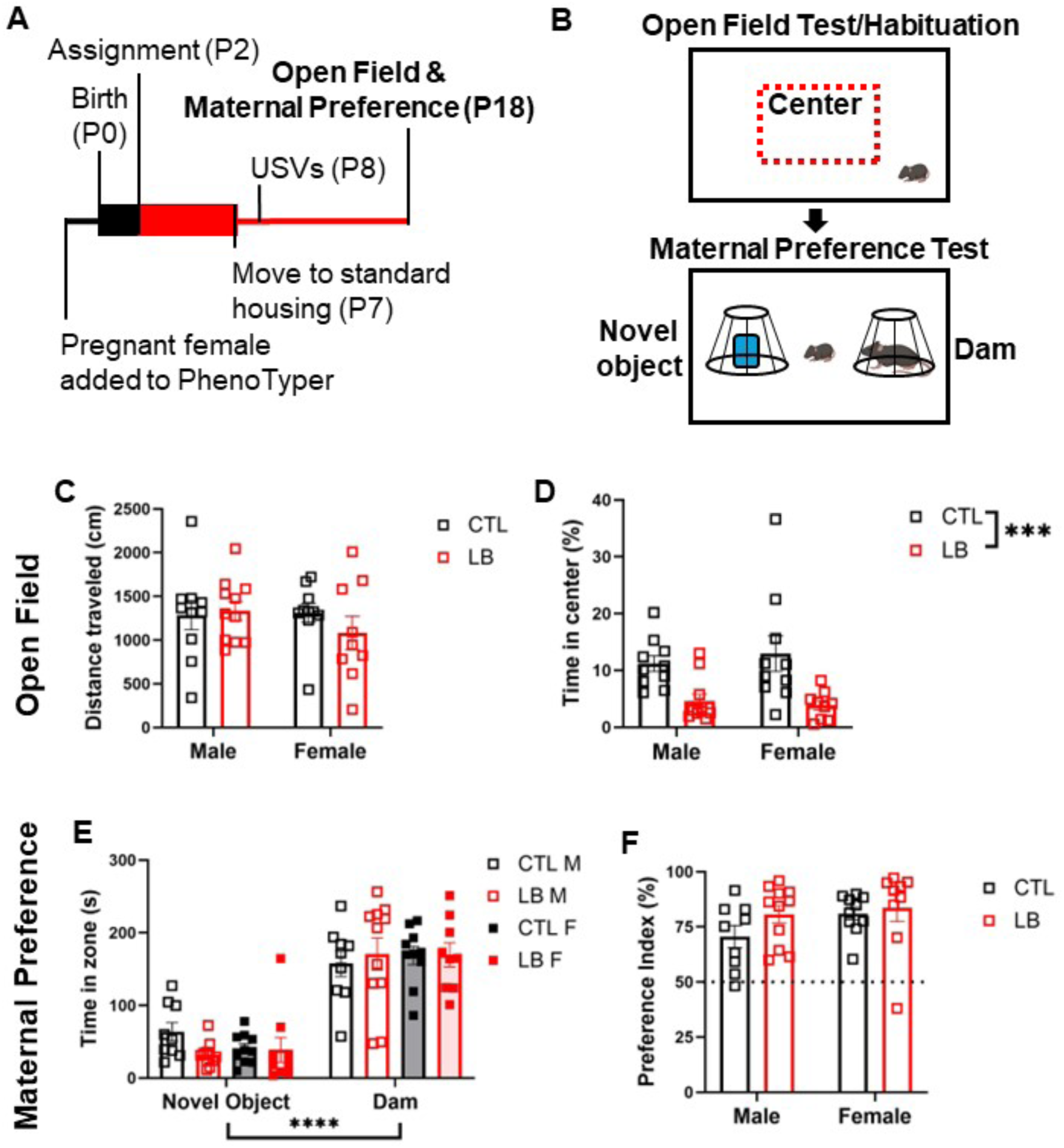
LB Pups Show More Anxiety-Like Behavior but Equivalent Maternal Preference at P18. **(A)** Experimental timeline. **(B)** Schematic of open field and maternal preference tests. **(C)** Distance moved during the open field test. Rearing, sex and interaction were all NS. **(D)** Percent time spent in the center of the arena. Rearing: F (1, 36) = 17.29, P = 0.0002; Sex: NS; Interaction: NS. **(E)** Time spent near the novel object and the dam. Zone: F (1, 70) = 137.4, P < 0.0001; rearing, sex, and all interactions were NS. **(F)** Preference index. Rearing, sex and interactions were all NS. N=5-6 litters and N=9-11 pups per group. 2 x 2 ANOVA in C, D, & F. 3-way rmANOVA in E. NS-non-significant, P > 0.05.

**Figure 7:**
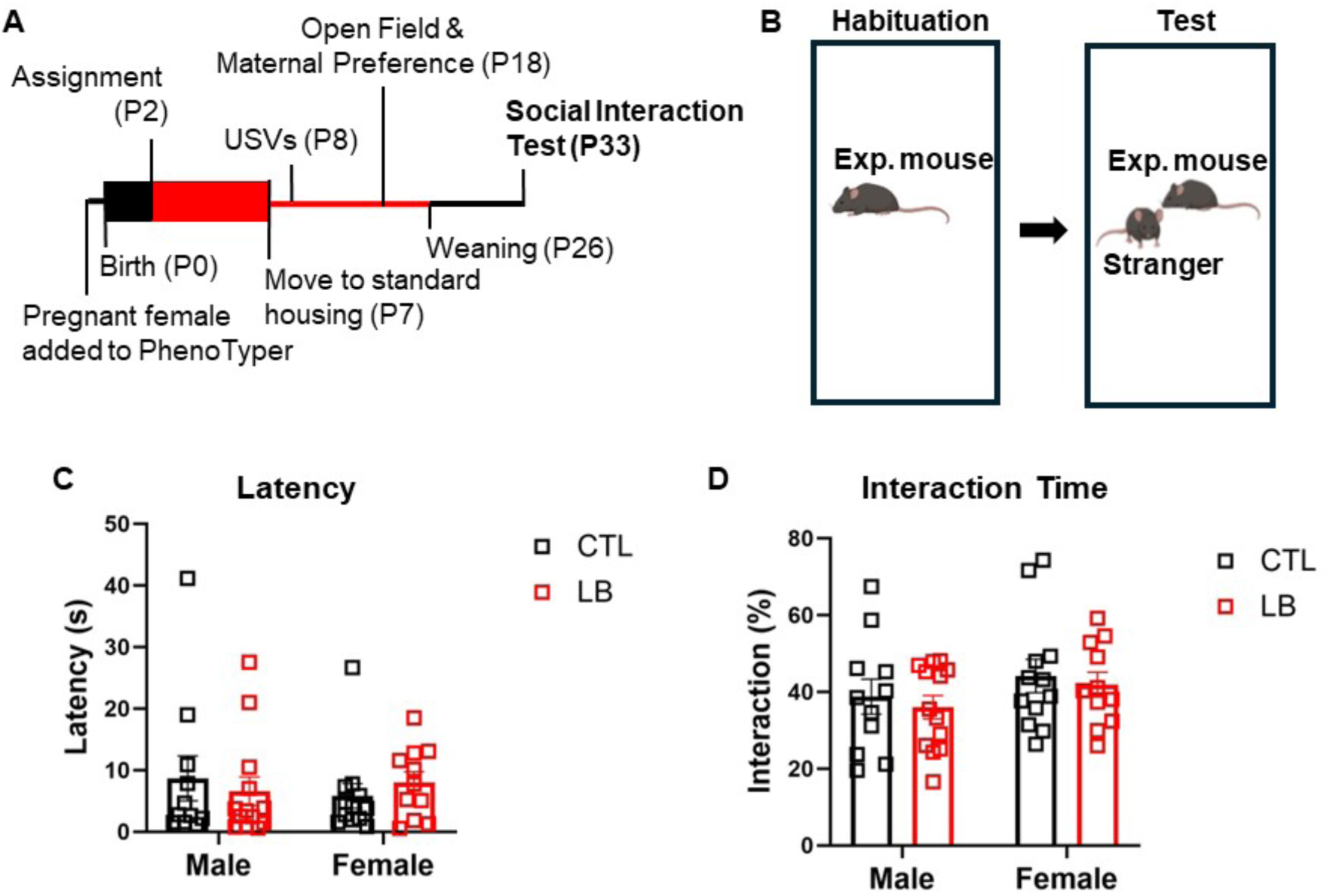
CTL and LB Pups Have Equivalent Social Interaction. **(A)** Timeline. **(B)** Test schematic. (**C**) Latency to first social contact initiated by the experimental mouse. Rearing, sex and interaction were all NS. **(D)** Cumulative time that the experimental mouse interacted with the stranger, expressed as a percent of test time. Rearing, sex and interaction were all NS. N = 6 litters and N = 11-13 pups per group. 2 x 2 ANOVA. NS-non-significant, P > 0.05.

## Results

### LB Increased Distance Travelled and Maternal Fragmentation

Total distance traveled, maternal fragmentation, and time in nest were calculated for each condition during the pre- and post-assignment periods across three cohorts (Fig 1A, n = 20-21 litters/condition). Cohort 1 (n = 6 litters/condition) was used for longitudinal behavior (P8, P18, & P33), cohort 2 (n = 9 litters/condition) was used for P7 corticosterone measurements, and cohort 3 (n = 6 CTL, 5 LB litters) was used for P13 behavior. Consistent with prior reports (Walker et al., 2017), LB dams exhibited a high degree of maternal fragmentation, characterized by frequent transitions in and out of the nest (Fig 1B, and supplemental videos S1A-CTL and S1B-LB). LB dams gathered all available bedding and nesting material to form a rudimentary nest; however, pups were often found scattered across the cage due to being dragged outside the nest while suckling (Fig. 1C, supplemental video S1C-LB).

A 2 x 2 repeated measures ANOVA (rmANOVA) found a significant interaction between pre- and post-assignment conditions and rearing conditions for hourly distance traveled, attributed to significantly greater distance traveled by LB dams relative to CTL dams during the post-assignment period (P2-P7; P < 0.0001), but not during the pre-assignment period (P0-P2, P = 0.14, Fig 1D). Comparisons in distance traveled between pre- and post-assignment found no significant changes in CTL dams (P = 0.39) as opposed to approximately 50% increase during post-assignment in LB dams (P<0.0001, Fig. 1D). To further examine the effects of rearing on hourly distance traveled during the post-assignment period, we conducted a three-way rmANOVA to characterize the effects of days and the dark-light phase as within-subjects variables and rearing as a between-subjects variable. This analysis revealed significant effects of day, rearing, the light-dark cycle, and a rearing by light-dark interaction (Fig 1E). The effect of day was due to a rapid decline in distance traveled from P2 to P3, as dams adjusted to the bedding change. From P3-7, both CTL and LB dams’ locomotor activity exhibited a distinct diurnal rhythm, with higher activity during the dark phase (active phase for mice) than the light phase (Fig 1E). This diurnal effect was greater for LB dams than CTL (Supplemental Fig. S1A).

A similar pattern was seen for maternal fragmentation, which is the frequency by which dams entered and exited the nest (Fig 1 F-G). Specifically, maternal fragmentation in CTL dams did not change after condition assignment (P = 0.49), whereas it was increased approximately threefold in LB dams (P < 0.0001, Fig. 1F). As with distance traveled, dams’ entrances/exits from the nest stabilized by P3, but was significantly higher in LB dams, especially during the dark phase (Fig. 1G and Fig. S1B). LB, but not CTL, dams, exhibited a diurnal rhythm in maternal fragmentation (light vs dark for CTL: P = 0.26, for LB: P < 0.0001; Fig. S1B).

Next, we evaluated the effects of pre-vs. post-assignment and rearing on time in the nest. Both CTL and LB dams decreased the average time spent on the nest from pre- to post-assignment; however, this reduction was greater for CTL than LB dams (CTL P < 0.0001, LB P = 0.0015), leading to LB dams spending more time than CTL on the nest from P2-P7 (CTL vs LB P = 0.004, Fig. 1H). Dams spent more time on the nest during the light (inactive) phase than during the dark (active) phase; as with movement and fragmentation, the difference between CTL and LB was greater during the dark phase (Supplemental Fig. S1C).

The effects of LB on dams’ movement and fragmentation were highly consistent across all three cohorts (Supplemental Fig. S2 A-B), underscoring the reproducible nature of these measurements. In contrast, the effect of rearing on time in nest was more variable and was significant for cohorts 1 and 2, but not in cohort 3 (Supplemental Fig. S2C). Finally, comparable intra-group variances were found between CTL and LB dams for distance traveled (Fig S3A) and time in the nest (Fig S3B), but not for maternal fragmentation for which inter-subject variability was significantly higher in LB dams (Fig S3C).

### Dam’s Locomotor Activity During the First Week of Life Negatively Correlated with Pup Bodyweight Throughout Development

A two-way ANOVA assessing the effects of rearing and sex on body weight at P8, P14, P18, P26, and P33 revealed significant and consistent lower body weight in LB pups compared to CTL from P8 to P33 (Fig 2A), with similar outcomes obtained in cohorts 2 & 3 (Fig S4). A transient, low-magnitude sex difference was noted at P14 but not at P18, with significant rearing x sex interactions detected at P26 and P33. These interactions were attributed to lower body weight in LB males, but not LB females, compared to their same-sex control groups (Fig 2A). Mean pup body weight for each litter was inversely correlated to the dam’s average movement from P2-P7 (Fig. 2B-F), a finding that was consistent with previous research utilizing standardized home cage and aluminum flooring (Rice et al., 2008; Treccani et al., 2021). This inverse correlation was strongest at P8 (R^2^ = 0.76, P < 0.001), but persisted at P14 (R^2^ = 0.52, P = 0.019, Fig. 2C) and showed a trend at P18 (R^2^ = 0.27, P = 0.08, Fig. 2D). Due to the significant interaction between rearing and body weight at P26 and P33, separate analyses were conducted for males and females at these ages. While both male and female pups’ weights had significant negative correlation with the dams’ movement at P26 (Fig. 2E), at P33 this relationship was only significant in males (Fig. 2F). Although the dams’ movement and maternal fragmentation positively correlated (R^2^ = 0.36, P = 0.039, N=12 litters), distance travelled provided a more reliable correlation with pup weight. These data reveal a strong negative and lasting relationship between the quality of maternal care during the first week of life, specifically distance traveled, and body weight that is more persistent in males.

### LB Increased Corticosterone Levels in P7 Pups

Previous work in mice has found a negative correlation between maternal fragmentation and corticosterone levels in P9 pups (Rice et al., 2008; Treccani et al., 2021) and similar findings in rats causally linked elevated corticosterone during the first week in life with abnormal attachment-like behavior in LB pups (Raineki et al., 2019; Opendak et al., 2020). Therefore, in a separate cohort of litters, we tested the dam and pups’ baseline blood corticosterone levels at P7 (Fig. 3A, for maternal behavior in this cohort see Fig S2). Corticosterone levels in LB pups were more variable and significantly higher in LB than CTL pups, with no differences between males and females (Fig. 3B). Dams’ corticosterone levels were comparable between CTL and LB conditions (Fig. 3C) with no significant correlation between corticosterone levels in the dam and her pups (CTL: r = 0.48, P = 0.34; LB: r = 0.62, P = 0.19; N = 6 dams per rearing condition, and 3-4 pups per litter). In agreement with previous reports (Rice et al., 2008; Treccani et al., 2021), we found a significant correlation between maternal fragmentation and corticosterone levels in P7 pups (Fig, 3D).

### LB Pups Had Reduced Vocalizations at P8 but Equivalent Maternal Buffering

A critical aspect of maternal attachment is the ability of the dam to reduce pups’ distress, a phenomenon known as “maternal buffering” (Bowlby, 1978; Kikusui et al., 2006; Gunnar et al., 2015). Here we assessed the effects of sex and rearing on maternal buffering in P8 pups (Fig. 4A, see also Fig. S2 for maternal behavior of cohort 1) by comparing the number of ultrasonic vocalizations (USVs) emitted by pups in response to a brief five-minute separation from the dam followed by a reunion with an anesthetized “aunt” (Fig 4B). We used anesthetized unrelated adult female instead of the dam in order to avoid exposing pups to anesthetic in the breast milk and because preliminary work from our lab and other groups has shown that at this age pups respond similarly to their dam and an unfamiliar aunt as long as they are fed the same diet (Opendak et al., 2020). A three-way rmANOVA assessing the effects of test phase (i.e. isolation vs with aunt), rearing, and sex demonstrated significant main effects of phase and rearing as well as significant interaction between phase and rearing and phase and sex (Fig 4C). Post-hoc analysis during the isolation phase revealed significant effect of rearing due to reduced vocalization in LB compared to CTL, with no significant effect of sex or interaction. No significant effects of rearing, sex or interaction between sex and rearing were found for the with aunt phase (Fig 4C). Body weight did not correlate with frequency of USVs during isolation (CTL: R^2^ = 0.0053, P = 0.67, LB: R^2^ = 0.017, P = 0.43; Fig. S5A), indicating the that reduced USVs observed in LB pups were not mediated by prematurity or low body weight. Furthermore, there was no significant correlation between maternal care and USVs during isolation in LB pups (distance traveled: r = -0.067, P = 0.89, fragmentation: r = -0.15, P = 0.78; Fig. S5B). Reunion with the aunt led to a robust buffering effect in all groups, with no significant effects of rearing, sex, or interaction (Fig 4D). Together, these data indicate that LB reduces distress vocalizations in response to maternal separation but does not disrupt maternal buffering.

### LB Impaired Proximity-Seeking Behavior in P13 Pups

Using a proximity-seeking approach toward an anesthetized dam as a measure of attachment-like behavior, Raineki et al. (2019) demonstrated that LB impairs approach behavior in P13 rat pups. These deficits were directly linked to elevated corticosterone levels in LB pups (Raineki et al., 2019). Given that LB increased corticosterone levels in P7 pups (Fig 3B), we tested whether similar deficits are observed in P13 LB mice pups and whether they correlated with abnormal maternal care (Fig 5 A-B and Fig S2 for maternal behavior in cohort 3). Ninety percent of all CTL pups (35/39) approached and maintained close proximity to the dam. In contrast, only 48% of LB pups (14/29) approached the dam (X^2^ (1, 14.2) = 3.77, P = 0.0002, Fig 5C), with no significant correlation between the proportion of pups that approached the dam within each litter and the dam’s average movement (CTL: r = -0.69, P = 0.13; LB: r = 0.54, P = 0.35). Similar results were obtained using the average distance maintained between the pup and the dam (P = 0.0001, Fig 5D). Within each LB litter, approximately half of the pups approached the dam while the remaining pups stayed in place or moved away from the dam (video S2), with no differences in weight, sex, or order of testing between LB pups that approached versus those that did not approach the dam (Fig. S6). For the LB pups that did approach the dam, there was a trend towards longer latency to approach (rearing: F(1, 45) = 3.91, P = 0.054, Fig 5E), but no difference in total interaction time (Fig. 5F). In contrast to findings by Raineki et al., CTL and LB pups did not differ in time spent on the dam’s dorsal or ventral side (F(1, 45) = 0.12, P = 0.73). In summary, LB reduces the likelihood of proximity-seeking behavior towards an anesthetized dam in P13 mice and rats. However, unlike findings in rats, LB did not alter the nature of the interaction in mice pups that engage with the dam, and the reduction in approach behavior was driven by notable individual differences within a litter.

### LB Pups Had Higher Anxiety-Like Behavior but Spent Equivalent Time with the Dam at P18

Attachment to a caregiver is thought to reduce anxiety and provide a secure base for exploring the environment. We therefore examined the effects of rearing and sex on exploratory behavior in P18 pups (Fig. 6A-B). CTL and LB pups were equally active exploring the empty arena (Fig. 6C), indicating the same degree of mobility despite LB pups’ lower body weight (Fig. 2A). However, LB pups spent approximately half the amount of time in the center arena as their CTL counterparts (Fig. 6D), indicating a higher level of anxiety-like behavior. Time in center did not correlate to the dams’ movement P2-7 (CTL: R^2^ = 0.05, P = 0.71; LB: R^2^ = 0.59, P = 0.13) and a similar difference in anxiety-like behavior was observed in CTL and LB mice raised in standardized cages instead of the PhenoTyper (Fig S7 A-B). Immediately following the open field test, maternal preference was tested by quantifying the time spent in the vicinity of the dam or a novel object placed under inverted wire cups in the same arena. All groups spent more time in the vicinity of the dam over the novel object, with no significant effects of rearing, sex or interaction (Fig. 6E). Maternal preference, calculated as the percent time spent in vicinity of dam over total time spent with either dam or the inanimate object, was significantly greater than 50% in all groups (P < 0.0001) indicating robust preference toward the dam with no significant effects of rearing, sex, or interaction (Fig. 6F). Similar findings were observed in an independent cohort raised in standard cages (FigS7 C-D). Thus, although LB pups have increased anxiety-like behavior at P18, they have normal maternal preference.

### No Effect of LB on Juvenile Social Interaction

According to attachment theory, deficits in attachment to a care giver also impact social affiliation later in life. Moreover, studies in both rats and mice have reported abnormal social interaction and play behavior in rodents raised under LB conditions that appear to be more prominent in males (Molet et al., 2016; Bolton et al., 2018; Birnie et al., 2023). Therefore, we tested social interaction at P33-34 with an age- and sex-matched CTL-raised mouse and analyzed interactions initiated by the experimental mouse (Fig. 7A-B). There were no differences between CTL and LB in latency to first interaction (Fig. 7C) or cumulative time spent interacting (Fig. 7D), nor did these metrics correlate to the dams’ movement from P2 to P7 (Latency: CTL: R^2^ = 0.0088, P = 0.86. LB: R^2^ = 0.13, P = 0.48. Total interaction time: CTL: R^2^ = 0.039, P = 0.71, LB: P = 0.31, R^2^ = 0.25).

## Discussion

Despite its conceptual and clinical promise, little is currently known about the underlying biology of attachment formation in humans, whether under normal or adverse conditions. However, the conserved nature of attachment behavior across diverse species, including rodents, suggests that rodent models may offer valuable insights (Sullivan and Opendak, 2021). Elegant work in rats has demonstrated that dams in poverty-like conditions, characterized by limited bedding and nesting (LB) exhibit erratic and abusive-like behaviors during the postnatal period (Walker et al., 2017). This abnormal form of maternal care is associated with elevated corticosterone levels in the pups (Rice et al., 2008; Raineki et al., 2019; Treccani et al., 2021), and the repeated association between maternal cues and corticosterone elevation leads to reduced cortical responsiveness to nurturing maternal care (Opendak et al., 2020), as well as abnormal amygdala development (Raineki et al., 2019). Furthermore, activation of the amygdala in response to maternal cues is directly responsible for deficits in attachment-like behavior observed in LB rat pups (Raineki et al., 2019).

The primary goals of this study were to determine whether LB induces similar attachment-like deficits in mice, where additional molecular tools are available to investigate the underlying biology. Furthermore, we employed a 24/7 automated home-cage monitoring system to better characterize maternal care and to correlate these changes with attachment-like deficits. To achieve this, we utilized the PhenoTyper system, which enables unbiased and automated quantification of maternal behavior throughout the light/dark cycle without investigator interference (Mingrone et al., 2020). Using three independent cohorts representing a total of 20-21 litters per condition, we confirmed that LB conditions consistently increased the total movement of dams by approximately 50% and the number of entries and exits of the nest (fragmentation) by 300% compared to dams in CTL conditions. These results confirm a substantial body of previous research indicating that LB increases maternal fragmentation (Walker et al., 2017). However, we show that these differences are more pronounced during the dark phase, underscoring the importance of measuring maternal behavior during both phases.

Average time spent on the nest decreased from pre- to post-assignment for both groups across all cohorts (Fig. 1H), suggesting that as pups mature, they require less maternal care. Overall, LB dams spent significantly less time on the nest, particularly during the dark phase (Fig. H-I). However, the relationship between rearing condition and time spent on the nest varied among the different cohorts. Specifically, LB dams spent more time on the nest in cohorts 1 and 2, but there was no significant difference between the groups in cohort 3 (Fig. S2). This variability may account for the conflicting outcomes reported in the literature, with some studies indicating increased time spent in the nest by LB dams (Gallo et al., 2019; Granata et al., 2021; Treccani et al., 2021), while others found no differences between the groups (Rice et al., 2008; Molet et al., 2016). LB dams may spend more time on the nest because their pups are smaller and/or as a compensatory mechanism for their increased erratic and fragmented care. Excessive time spent on the nest may also elevate the rate of rough handling and stepping on pups, as documented in previous studies (Gallo et al., 2019; Raineki et al., 2019). Collectively, time spent in the nest is a more variable aspect of the LB paradigm that requires further characterization and assessment of its impact on attachment-like behaviors.

Pup weight at P8 was strongly and inversely correlated with the dams’ movement from P2-P7. This finding is consistent with previous work (Rice et al., 2008), but the magnitude of the correlation was higher when using continuous monitoring (R^2^ = -0.76, Fig 2B vs. R^2^ = -0.26 in Rice et al., 2008). Furthermore, the impact of the dam’s movement from P2-7 on body weight remained significant at P14, P18, and P26 for both sexes. This relationship continued to be significant for males but not for females at P33. These findings reveal a novel relationship between erratic maternal care early in life and long-term growth, which appears to be more persistent in male offspring. The mechanisms responsible for the reduced body weight observed in LB mice have yet to be clarified; however, (Noviana et al., 2023) found that interventions aimed at increasing secure attachment were effective in reducing stunting in high-risk populations, demonstrating a potentially significant link between abnormal attachment and deficits in normal growth. We also replicated previous work showing that LB increases corticosterone levels in pups in a manner that correlated with maternal fragmentation (Rice et al., 2008; Raineki et al., 2019; Treccani et al., 2021), but found no differences in corticosterone levels between CTL and LB dams.

USVs at P8 are the murine equivalent of a baby’s cry; they serve as distress calls designed to elicit caregiving (Zimmer et al., 2019). Our findings indicate that P8 LB pups produced fewer USVs during isolation compared to CTL pups. This reduction was not correlated with decreased body weight or maternal fragmentation. A similar decrease in USVs was reported previously but only in female LB pups (Granata et al., 2021). Given that Agouti-related peptide (AgRP)-expressing neurons in the arcuate nucleus have been found to promote isolation-related USVs in mouse pups (Zimmer et al., 2019), it would be interesting to know whether LB inhibits AgRP activation in response to isolation. Since LB pups are accustomed to a poor nest environment and lower-quality maternal care, they may experience less distress during isolation than CTL pups. Alternatively, they may be less motivated to elicit retrieval by the dam due to her frequent departures from the nest and/or rough handling within the nest. Nevertheless, reunion with an aunt reduced USVs by approximately 70% across all groups. Together, the reduced USVs observed in response to separation in eight-day-old LB pups suggest an avoidant-like attachment style that is also highly responsive to maternal buffering.

Reduced rates of maternal approach observed at P13 among LB pups are consistent with findings in rats (Raineki et al., 2019). However, there are notable differences between our results and those reported in rats. For instance, Raineki and colleagues identified deficits in nipple attachment and relocation to the back of the dam in LB pups, whereas we primarily observed deficits in approach behavior. Furthermore, we noted significant individual variability in approach behavior within LB litters. These individual differences were not attributable to differences in weight, sex, or testing order. Additional studies are necessary to elucidate the underlying biology that drives these individual differences and to determine whether they represent stable long-term changes in other behavioral tests. The reduced rate of proximity-seeking in 13-day-old pups further supports the notion that LB causes avoidant-like attachment deficits.

In the open field test, 18-day-old LB pups spent less time exploring the center, which is consistent with higher rates of anxiety observed in children with insecure attachment (Feeney, 2000; Hornor, 2019; Gregory et al., 2020). Both CTL and LB pups showed a strong preference for the dam over an inanimate object in the maternal preference test, indicating normal attachment-like behavior in this test. Similar results were obtained in CTL and LB litters raised under standard conditions (i.e., not in the PhenoTyper), indicating that these outcomes are robust and do not require specialized housing conditions. Social exploration was comparable in CTL and LB P33 juvenile mice during free interactions with a CTL-reared age-and sex-matched stranger, indicating that some attachment-like behaviors are resilient even in mice exposed to high levels of maternal fragmentation.

In conclusion, automated continuous home-cage monitoring provides a powerful method for quantifying maternal care in mice raised under CTL and LB conditions. Using this approach, we extended previous observations by demonstrating that differences in maternal behavior are more pronounced during the dark phase and that time spent on the nest is a more variable outcome. Maternal fragmentation exhibited a strong negative correlation with body weight and resulted in stunted growth that was more persistent in males. LB pups displayed avoidant-like attachment deficits, as evidenced by reduced USVs in response to maternal separation at P8, deficits in approaching an anesthetized dam at P13, and decreased exploration of the center in the open-field test at P18. None of these deficits were correlated with maternal fragmentation, suggesting a possible threshold effect rather than a graded effect. This hypothesis is keeping with the notion of the “good enough” parent, proposed by Donald Winnicott in the 1950s (Taylor et al., 2009). Intriguing within-litter individual differences in approach behavior were observed in both CTL and LB groups, with 10% of CTL pups versus 50% of LB pups exhibiting non-approach behavior. No deficits were observed in the maternal preference test at P18 or in social exploration at P33, underscoring the robustness of these attachment-like behaviors despite severe perturbations in maternal care. Collectively, these findings reveal important similarities and differences in the effects of LB on attachment-like behavior in rats and mice and lay the groundwork for further investigation into the neurobiological underpinnings of attachment in mice.

## Acknowledgements.

This work was supported by: NIMH R01MH130825 (MOD, AK), NIMH R01MH136490 (AK), the Clinical Neuroscience Division of the VA National Center for PTSD (AK), JMM was funded by FPU21/01318 (MICIU/AEI, Spain), Movilidad MOV24-16 and MOV25-06 (IBIMA Plataforma Bionand, Málaga, Spain), and Acción 321 PPID Universidad de Málaga-Banco Santander (Spain).

## Supplemental Figures

MacDowell Kaswan Z., et al. (2025)

**Figure S1:**
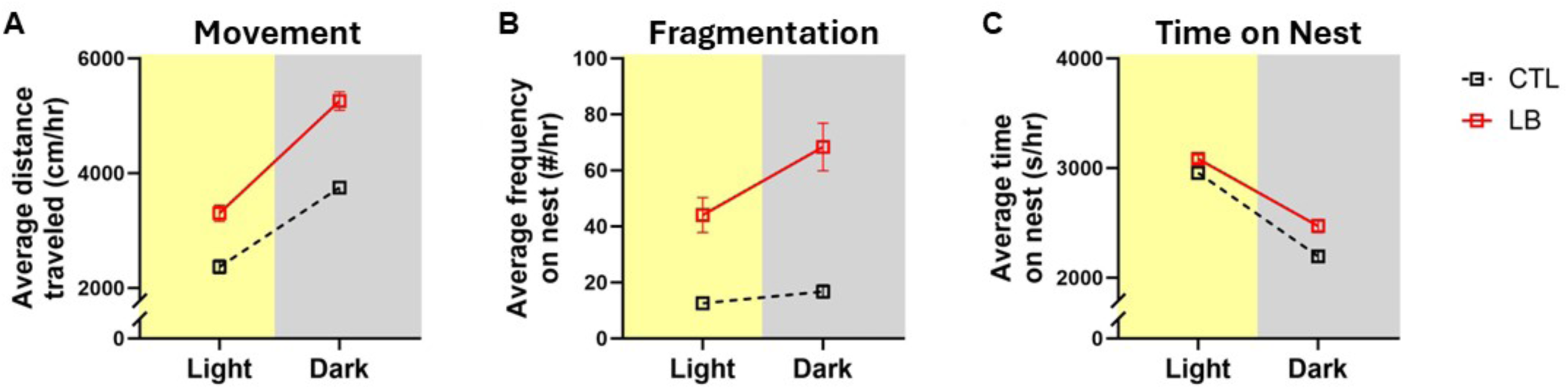
Differences in Maternal Behavior Are More Prominent During the Dark Phase. Maternal behavior is shown for the light and dark phases of P3-6. **(A)** Average hourly distance travelled. Rearing: F(1, 39) = 62.72, P < 0.0001; Light/dark: F(1, 39) = 266.8, P < 0.0001; Subject: F(39, 39) = 2.28, P = 0.0058; Rearing x Light/dark interaction: F(1, 39) = 7.96, P = 0.0075. Post-hoc analysis: CTL vs LB (light phase), CTL vs LB (dark phase), Light vs Dark (CTL), and Light vs Dark (LB) all P < 0.0001. **(B)** Average hourly frequency on the nest. Rearing: F(1, 39) = 34.28, P < 0.0001; Light/dark: F(1, 39) = 28.61, P < 0.0001; Subject: F(39, 39) = 7.07, P < 0.0001; Rearing x Light/dark interaction: F(1, 39) = 14.14, P = 0.0006. Post-hoc analysis: CTL vs LB (light phase) P < 0.0001, CTL vs LB (dark phase) P < 0.0001, Light vs Dark (CTL) P = 0.26, Light vs Dark (LB) P < 0.0001. **(C)** Average hourly time on the nest. Rearing: F(1, 39) = 14.66, P = 0.0005; Light/dark: F(1, 39) = 446.2, P < 0.0001; Subject: F(39, 39) = 2.63, P = 0.0016; Rearing x Light/dark interaction: F(1, 39) = 5.22, P = 0.028. Post-hoc analysis: CTL vs LB (light phase) P = 0.43, CTL vs LB (dark phase) P < 0.0001, Light vs Dark (CTL) P < 0.0001, Light vs Dark (LB) P < 0.0001. N=21 litters per rearing condition. Analyzed by 2 x 2 rmANOVA with post-hoc uncorrected Fisher’s least significant difference test.

**Figure S2.**
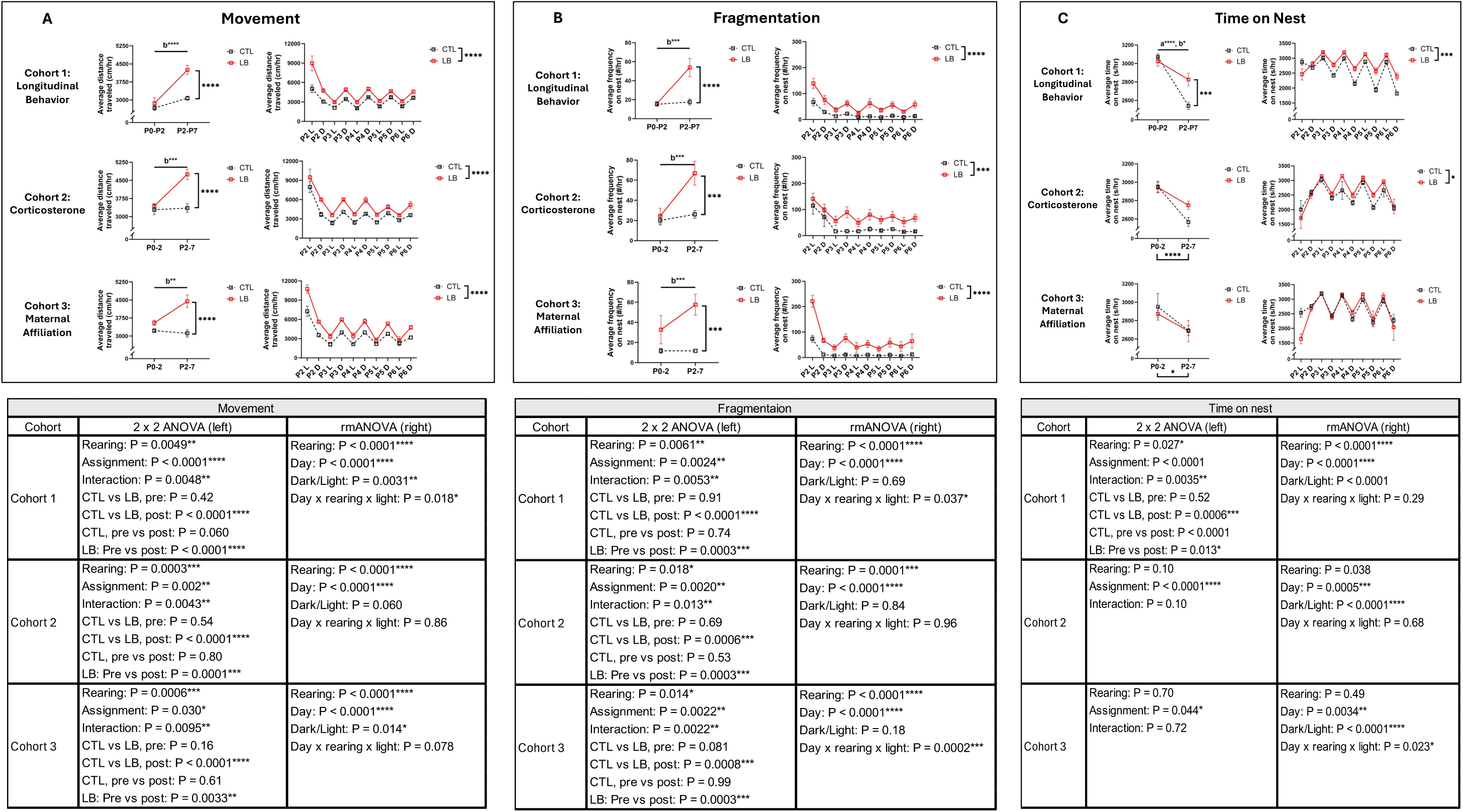
Reproducible and Robust Effects of Rearing Observed for Distance Traveled and Maternal Fragmentation, But Not for Time on Nest. Effects on maternal movement (**A**), fragmentation (**B**), and time on nest (**C**) for the three cohorts tested. Cohort 1 (n = 6 litters/condition) was used for longitudinal behavior (P8, P18, & P33), cohort 2 (n = 9 litters/condition) was used for P7 corticosterone measurements, and cohort 3 (n = 6 CTL, 5 LB litters) was used for P13 maternal affiliation behavior.

**Figure S3:**
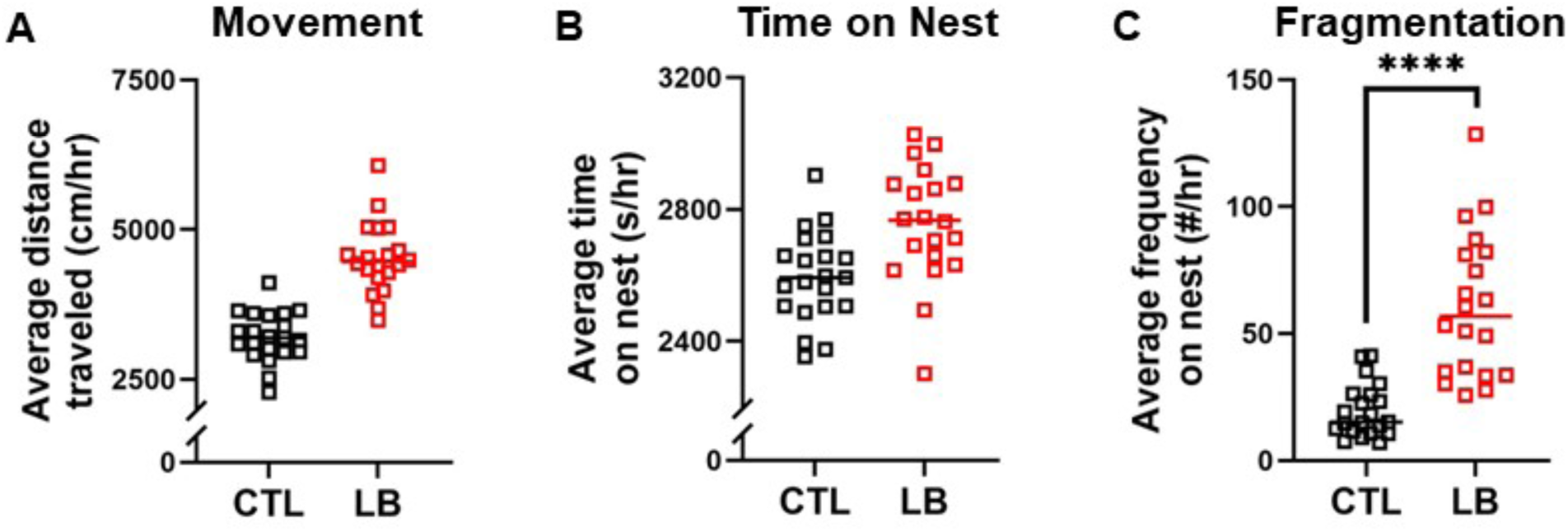
Effects of Rearing on Variance in Dam’s Behavior from P2 to P7. **(A)** Variance in movement, F (19,20) = 2.20, P = 0.13. **(B)** Variance in time on nest, F (19,20) = 1.69, P = 0.25. (C) Variance in fragmentation, F (19,20) = 7.47, P < 0.0001. N = 20-21 litters per rearing condition.

**Figure S4:**
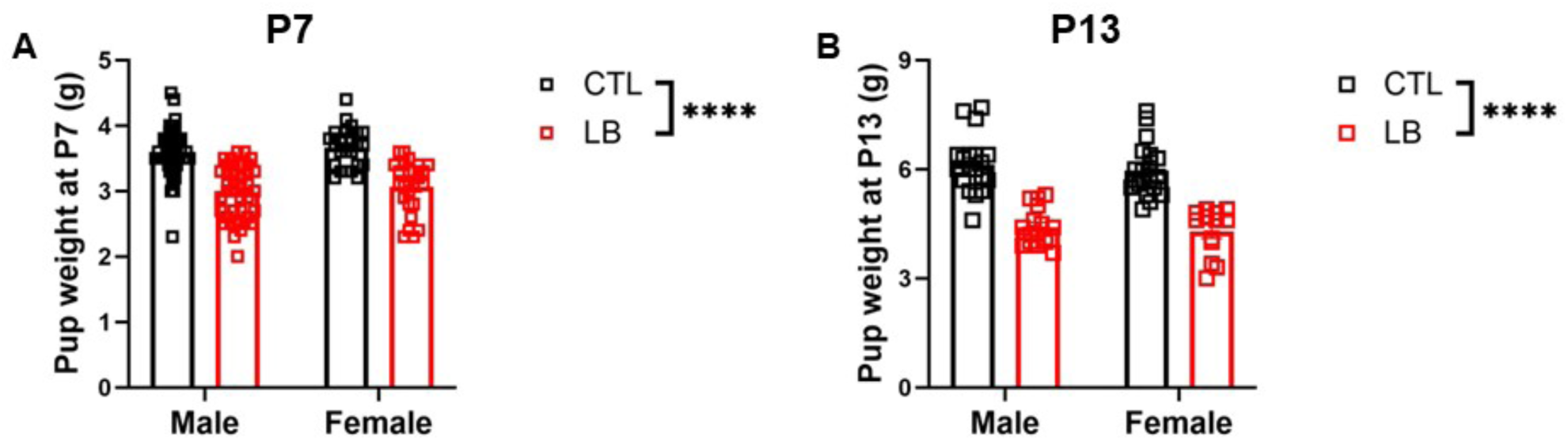
Pup Weights at P7 and P13. Pup weights from cohorts 2 **(A)** and 3 **(B)**. **(A)** LB pups weigh significantly less than CTL pups at P7 without an effect of sex. N=24-40 per group, from N=9 litters per rearing condition. Rearing: F(1, 124) = 74.84, P < 0.0001; Sex: F(1, 124) = 1.227, P = 0.27; Interaction: F(1, 124) = 0.0046, P = 0.95. **(B)** LB pups weigh significantly less than CTL pups at P13 without an effect of sex. N=13-20 per group, from N = 6 CTL and 5 LB litters. Rearing: F(1, 64) = 101.9, P < 0.0001; Sex: F(1, 64) = 0.79, P = 0.38; Interaction: F(1, 64) = 0.017, P = 0.90. Analysis by 2-way ANOVA.

**Figure S5:**
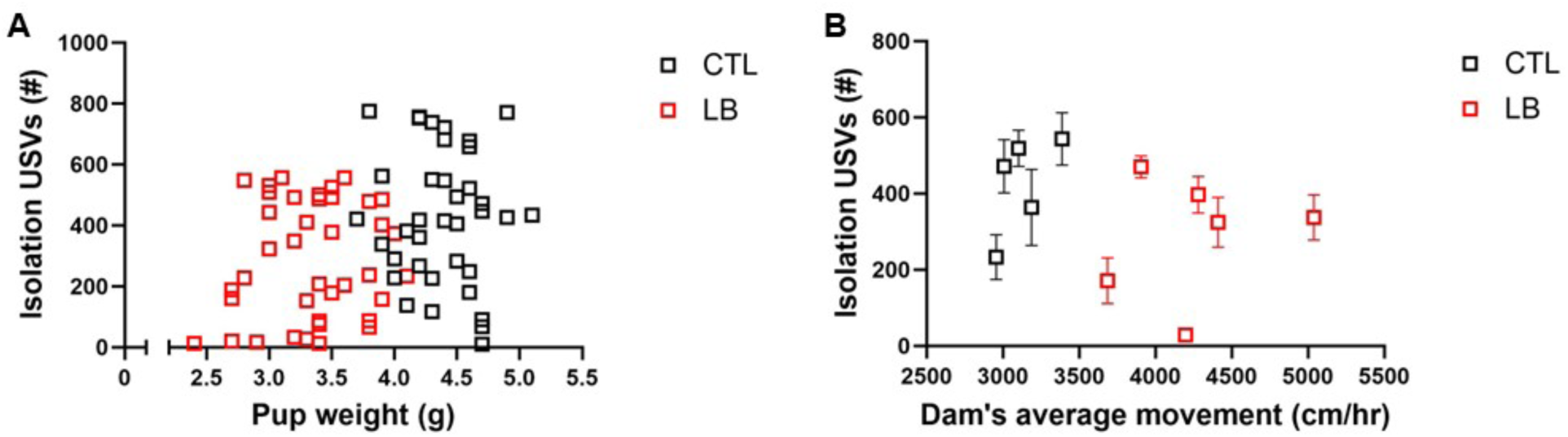
P8 Isolation USVs Do Not Correlate to Pup Weight or Dam Behavior. **(A)** Correlation between P8 CTL and LB pup weights and number of USVs during the isolation phase of the maternal buffering test. N = 37-39 per condition, sexes are combined for analysis since neither behavior nor weight varied by sex. CTL: R^2^ = 0.0053, P = 0.67; LB: R^2^ = 0.017, P = 0.43. **(B)** Dams’ average movement P2-7 does not correlate with pups’ USVs during isolation at P8. CTL: R^2^ = 0.36, P = 0.29; LB: R^2^ = 0.03, P = 0.74.

**Figure S6:**
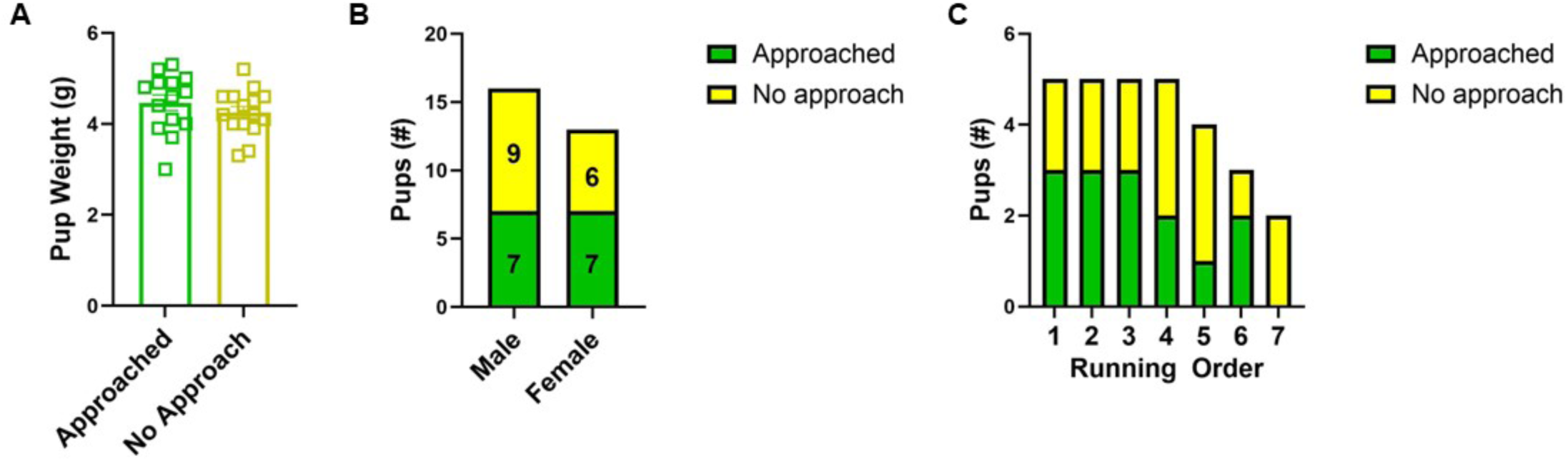
Effects of Body Weight, Sex, and Order of Testing on Approach behavior in P13 LB pups. **(A)** Body weight, t(27) = 0.98, P = 0.34. **(B)** Sex, Fisher’s exact test, P = 0.72. **(E)** Testing order. Fisher’s exact test, P = 0.83. N = 29 pups from N = 5 litters, sexes combined for A & C.

**Figure S7:**
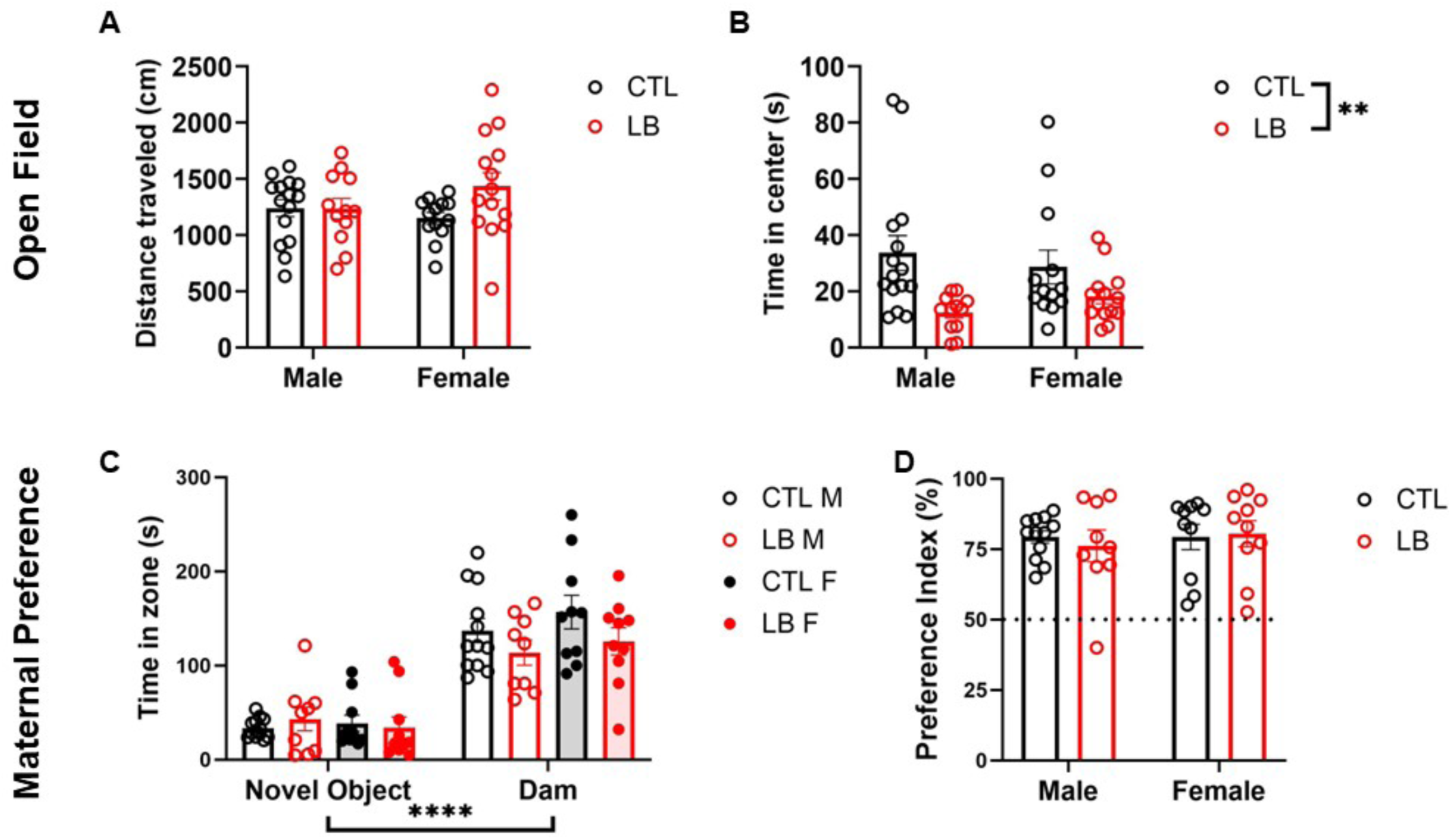
Similar P18 Results in a Cohort of Standard-housed Pups. **(A)** Distance moved during the open field test. Rearing: F (1, 50) = 2.39,P = 0.13; Sex: F (1, 50) = 0.37, P = 0.55; Interaction: F (1, 50) = 2.56, P = 0.12. **(B)** Time spent in the center of the arena. Rearing: F (1, 50) = 11.10, P = 0.0016; Sex: F (1, 50) = 0.0076, P = 0.93; Interaction: F (1, 50) = 1.25, P = 0.27. **(C)** Time spent near the novel object and the dam. Zone: F (1, 37) = 116.5, P < 0.0001; Rearing: F (1, 37) = 2.19, P = 0.15; Sex: F(1, 37) = 0.65, P = 0.42; Zone x Sex: F(1, 37) = 0.1, P =0. 32; Zone x Rearing: F (1, 37) = 2.81, P = 0.10; Sex x Rearing: F (1, 37) = 0.41, P = 0.52; Zone x Sex x Rearing: F (1, 37) = 0.033, P = 0.86. **(D)** Preference index. Rearing: F (1, 37) = 0.057, P = 0.81; Sex: F (1, 37) = 0.26, P = 0.62; Interaction: F (1, 37) = 0.26, P = 0.61. N = 9-12 pups per group from N=4-5 litters. 2 x 2 ANOVA in A, B, & D. 3-way rmANOVA in C.

## References

1. Ahmed S, Polis B, Jamwal S, Sanganahalli BG, MacDowell Kaswan Z, Islam R, Kim D, Bowers C, Giuliano L, Biederer T, Hyder F, Kaffman A (2024) Transient impairment in microglial function causes sex-specific deficits in synaptic maturity and hippocampal function in mice exposed to early adversity. Brain, behavior, and immunity 122:95–109.

2. Birnie MT, Short AK, de Carvalho GB, Taniguchi L, Gunn BG, Pham AL, Itoga CA, Xu X, Chen LY, Mahler SV, Chen Y, Baram TZ (2023) Stress-induced plasticity of a CRH/GABA projection disrupts reward behaviors in mice. Nature communications 14:1088.

3. Bolton JL, Molet J, Regev L, Chen Y, Rismanchi N, Haddad E, Yang DZ, Obenaus A, Baram TZ (2018) Anhedonia Following Early-Life Adversity Involves Aberrant Interaction of Reward and Anxiety Circuits and Is Reversed by Partial Silencing of Amygdala Corticotropin-Releasing Hormone Gene. Biol Psychiatry 83:137–147.

4. Bowlby J (1978) Attachment theory and its therapeutic implications. Adolesc Psychiatry 6:5–33.

5. Diamond GS, Kobak RR, Krauthamer Ewing ES, Levy SA, Herres JL, Russon JM, Gallop RJ (2019) A Randomized Controlled Trial: Attachment-Based Family and Nondirective Supportive Treatments for Youth Who Are Suicidal. J Am Acad Child Adolesc Psychiatry 58:721–731.

6. Feeney JA (2000) Implications of attachment style for patterns of health and illness. Child Care Health Dev 26:277–288.

7. Fonseca AH, Santana GM, Bosque Ortiz GM, Bampi S, Dietrich MO (2021) Analysis of ultrasonic vocalizations from mice using computer vision and machine learning. Elife 10.

8. Gallo M, Shleifer DG, Godoy LD, Ofray D, Olaniyan A, Campbell T, Bath KG (2019) Limited Bedding and Nesting Induces Maternal Behavior Resembling Both Hypervigilance and Abuse. Frontiers in behavioral neuroscience 13:167.

9. Granata L, Valentine A, Hirsch JL, Honeycutt J, Brenhouse H (2021) Trajectories of Mother-Infant Communication: An Experiential Measure of the Impacts of Early Life Adversity. Frontiers in human neuroscience 15:632702.

10. Gregory M, Dymand LK, Sharman R (2020) A review of attachment-based parenting interventions: Recent advances and future considerations. Australian Journal of Psychology 72:109–122.

11. Gunnar MR, Hostinar CE, Sanchez MM, Tottenham N, Sullivan RM (2015) Parental buffering of fear and stress neurobiology: Reviewing parallels across rodent, monkey, and human models. Soc Neurosci 10:474–478.

12. Hepworth AD, Berlin LJ, Martoccio TL, Cannon EN, Berger RH, Harden BJ (2020) Supporting Infant Emotion Regulation Through Attachment-Based Intervention: a Randomized Controlled Trial. Prev Sci 21:702–713.

13. Hornor G (2019) Attachment Disorders. J Pediatr Health Care 33:612–622.

14. Islam R, White DR, Arefin TM, Mehta S, Liu X, Polis B, Giuliano L, Ahmed S, Bowers C, Zhang J, Kaffman A (2024) Early Adversity Causes Sex-Specific Deficits in Perforant Pathway Connectivity and Contextual Memory in Adolescent Mice. Biology of sex differences 15.

15. Izaki A, Verbeke W, Vrticka P, Ein-Dor T (2024) A narrative on the neurobiological roots of attachment-system functioning. Commun Psychol 2:96.

16. Kikusui T, Winslow JT, Mori Y (2006) Social buffering: relief from stress and anxiety. Philos Trans R Soc Lond B Biol Sci 361:2215–2228.

17. Londono Tobon A, Condon E, Slade A, Holland ML, Mayes LC, Sadler LS (2023) Participation in an Attachment-Based Home Visiting Program Is Associated with Lower Child Salivary C-Reactive Protein Levels at Follow-Up. J Dev Behav Pediatr 44:e292–e299.

18. Mingrone A, Kaffman A, Kaffman A (2020) The Promise of Automated Home-Cage Monitoring in Improving Translational Utility of Psychiatric Research in Rodents. Frontiers in neuroscience 14:618593.

19. Molet J, Heins K, Zhuo X, Mei YT, Regev L, Baram TZ, Stern H (2016) Fragmentation and high entropy of neonatal experience predict adolescent emotional outcome. Translational psychiatry 6:e702.

20. Noviana U, Ekawati H, Mufarika M, Hasinuddin, Haris M (2023) Mother’s Behavior Attachment Model in Care for Stunting Prevention in Bangkalan District. Journal Of Nursing Practice 7:57–66.

21. Opendak M, Theisen E, Blomkvist A, Hollis K, Lind T, Sarro E, Lundstrom JN, Tottenham N, Dozier M, Wilson DA, Sullivan RM (2020) Adverse caregiving in infancy blunts neural processing of the mother. Nature communications 11:1119.

22. Raineki C, Opendak M, Sarro E, Showler A, Bui K, McEwen BS, Wilson DA, Sullivan RM (2019) During infant maltreatment, stress targets hippocampus, but stress with mother present targets amygdala and social behavior. Proc Natl Acad Sci U S A 116:22821–22832.

23. Rice CJ, Sandman CA, Lenjavi MR, Baram TZ (2008) A novel mouse model for acute and long-lasting consequences of early life stress. Endocrinology 149:4892–4900.

24. Ross KM, Cole S, Sanghera H, Anis L, Hart M, Letourneau N (2021) The ATTACH program and immune cell gene expression profiles in mothers and children: A pilot randomized controlled trial. Brain Behav Immun Health 18:100358.

25. Sullivan RM, Opendak M (2021) Neurobiology of Infant Fear and Anxiety: Impacts of Delayed Amygdala Development and Attachment Figure Quality. Biol Psychiatry 89:641–650.

26. Taylor J, Lauder W, Moy M, Corlett J (2009) Practitioner assessments of ’good enough’ parenting: factorial survey. J Clin Nurs 18:1180–1189.

27. Treccani G, Yigit H, Lingner T, Schleubetaner V, Mey F, van der Kooij MA, Wennstrom M, Herzog DP, Linke M, Fricke M, Schmeisser MJ, Wegener G, Mittmann T, Trotter J, Muller MB (2021) Early life adversity targets the transcriptional signature of hippocampal NG2+ glia and affects voltage gated sodium (Na(v)) channels properties. Neurobiology of stress 15:100338.

28. Walker CD, Bath KG, Joels M, Korosi A, Larauche M, Lucassen PJ, Morris MJ, Raineki C, Roth TL, Sullivan RM, Tache Y, Baram TZ (2017) Chronic early life stress induced by limited bedding and nesting (LBN) material in rodents: critical considerations of methodology, outcomes and translational potential. Stress 20:421–448.

29. Wright B, Edginton E (2016) Evidence-Based Parenting Interventions to Promote Secure Attachment: Findings From a Systematic Review and Meta-Analysis. Glob Pediatr Health 3:2333794X16661888.

30. Zimmer MR, Fonseca AHO, Iyilikci O, Pra RD, Dietrich MO (2019) Functional Ontogeny of Hypothalamic Agrp Neurons in Neonatal Mouse Behaviors. Cell 178:44–59 e47.

